# Fibroblast growth factor is predicted to dominate MAPK activation by pro-angiogenic factors

**DOI:** 10.1101/368415

**Authors:** Min Song, Stacey D. Finley

## Abstract

Angiogenesis is important in physiological and pathological conditions, as blood vessels provide nutrients and oxygen needed for tissue growth and survival. Therefore, targeting angiogenesis is a prominent strategy in both tissue engineering and cancer treatment. However, not all of the approaches to promote or inhibit angiogenesis lead to successful outcomes. Angiogenesis-based therapies primarily target pro-angiogenic factors such as vascular endothelial growth factor-A (VEGF) or fibroblast growth factor (FGF) in isolation, and there is a limited understanding of how these promoters combine together to stimulate angiogenesis. Thus, more quantitative insight is needed to understand their interactions. In this study, we have trained and validated a detailed mathematical model to quantitatively characterize the crosstalk of FGF and VEGF intracellular signaling. The model focuses on FGF- and VEGF-induced mitogen-activated protein kinase (MAPK) signaling and phosphorylation of extracellular regulated kinase (ERK), which promote cell proliferation. We apply the model to predict the dynamics of phosphorylated ERK (pERK) in response to the stimulation by FGF and VEGF individually and in combination. The model predicts that FGF plays a dominant role in promoting ERK phosphorylation, compared to VEGF. The modeling predictions show that VEGFR2 density and trafficking parameters significantly influence the level of VEGF-induced pERK. The model matches experimental data and is a framework to synthesize and quantitatively explain experimental studies. Ultimately, the model provides mechanistic insight into FGF and VEGF interactions needed to identify potential targets for pro-or anti-angiogenic therapies.

## Introduction

Angiogenesis is the formation of new blood capillaries from pre-existing blood vessels. The essential role of blood vessels in delivering nutrients makes angiogenesis important in the survival of tissues, including tumor growth. Angiogenesis also provides a route for tumor metastasis. Thus, targeting angiogenesis is a prominent strategy in many contexts, for example, in both tissue engineering and cancer treatment.

In the context of tissue engineering, there is a large demand for organs needed for transplant surgery, but a great shortage of donors. In 2015, the US Department of Health and Human Services estimated that there were more than 120,000 patients on waiting lists for transplant surgery. Due to the limited number of donor organs, researchers have looked to engineered tissues as substitutes to improve or replace damaged tissues. One requirement for successful engraftment of synthetic tissues is to ensure sufficient transport of nutrients and gases such as oxygen to the cells [1]. The principal mode of transport for small molecules, including oxygen, is passive diffusion. However, oxygen diffusion is limited to less than approximately 200 µm [1-3], while most synthetic tissues are thicker than 1 cm [1]. Therefore, the long-term viability of engineered tissue constructs depends on growth of new vessels from host tissue, and stimulating new blood vessel formation is an important strategy for tissue engineering [3].

Alternatively, the formation of new blood vessels is important for cancer growth and metastasis. Thus, inhibiting angiogenesis is a strategy for cancer treatment. There are two major approaches for these “anti-angiogenic” strategies: blocking pro-angiogenic intracellular signaling directly and targeting extracellular ligands that promote angiogenesis [4]. For example, bevacizumab, an anti-angiogenic agent was designed to bind to the pro-angiogenic protein vascular endothelial growth factor (VEGF) extracellularly, preventing VEGF from binding to its receptors and inhibiting VEGF-mediated signaling [5, 6].

Unfortunately, not all approaches to promote or inhibit angiogenesis lead to successful outcomes. For example, clinical trials have shown no effective improvement in FGF-induced [7] or VEGF-induced [8] angiogenesis. Also, bevacizumab has limited effects in certain cancer types, and it is no longer approved for the treatment of metastatic breast cancer due to its disappointing results [9]. Thus, there is a need to better understand the molecular interactions and signaling required for new blood vessel formation, in order to establish more effective therapeutic strategies.

The established angiogenesis-based therapies primarily target pro-angiogenic factors such as FGF and VEGF in isolation. However, both FGF and VEGF bind to their receptors to initiate mitogen-activated protein kinase (MAPK) signaling and phosphorylate ERK, the final output of the MAPK pathway [10, 11]. This signaling pathway promotes cell proliferation in the early stages of angiogenesis. Additionally, the combined effects of FGF and VEGF have been shown to be greater than their individual effects [12, 13]. A quantitative understanding of how these promoters combine together to stimulate angiogenesis could greatly benefit the current pro-and anti-angiogenic therapies.

Mathematical modeling is a useful tool to predict the molecular response mediated by angiogenic factors. For example, Mac Gabhann and Popel studied interactions between VEGF isoforms, VEGF receptors (VEGFR1, VEGFR2, NRP1), and the extracellular matrix using a molecular-detailed model. The model predicted that blocking Neuropilin-VEGFR coupling is more effective in reducing VEGF-VEGFR2 signaling than blocking Neuropilin-1 expression or binding of VEGF to Neuropilin-1 [14]. Stefanini *et al*. constructed a pharmacokinetic model that studied VEGF distribution after intravenous administration of bevacizumab, and they found that plasma VEGF was increased after treatment [15]. Filion and Popel explored myocardial deposition and retention of FGF after intracoronary administration of FGF using a computational model. The model predicted that the response time is dependent on the reaction time of the binding of FGF to FGFR rather than the FGF diffusion time. Receptor secretion and internalization have also been predicted to be crucial in FGF dynamics [16]. Wu and Finley characterized the intracellular signaling of TSP1-induced apoptosis and predicted response of cell population to TSP1-mediated apoptosis by mathematical modeling [17]. Zheng *et al*. integrated the effects of VEGF, angiopoietins (Ang1 and Ang2) and platelet-derived growth factor-B (PDGF-B) on endothelial proliferation, migration, and maturation using mathematical modeling. Their model illustrated that competition between Ang1 and Ang 2 acts as an angiogenic switch and that combining anti-pericyte and anti-VEGF therapy is more effective than anti-VEGF therapy alone in inducing blood vessel regression [18].

In the present study, we aim to quantitatively characterize the crosstalk between FGF and VEGF in MAPK signaling leading to phosphorylated ERK (pERK). We focus on this species because pERK promotes cell proliferation [19] and is mostly found in active, rather than quiescent, endothelial cells [20]. We constructed a computational model that incorporates the molecular interactions between FGF, VEGF, and their receptors, leading to MAPK signaling. We apply the model to explore how FGF and VEGF promote ERK phosphorylation. To the best of our knowledge, this is the first model that studies FGF and VEGF interactions together on a molecular level. Our model predicts the combination effects of FGF and VEGF stimulation and shows that FGF plays a dominant role in promoting ERK phosphorylation. Using this model, we also investigated the effects of the VEGF receptor VEGFR2, including how VEGFR2 density and trafficking parameters influence the ERK response. Predicting the effects of these quantities helps identify potential targets for enhancing pro-or anti-angiogenic therapies. More broadly, this model provides a framework to study the efficacy of angiogenesis-based therapies.

## Results

### The mathematical model captures the main features of FGF-and VEGF-stimulated ERK phosphorylation dynamics

We constructed a computational model that characterizes FGF and VEGF interactions leading to ERK phosphorylation (Figure 1). Signaling is mediated by FGF and VEGF binding to their respective receptors, leading to pERK. The model was trained against published experimental measurements [21-23] using Particle Swarm Optimization (PSO) for parameter estimation. We note that pERK response stimulated by FGF was measured using the non-small cell lung cancer cell line NCI-H1730 [21], while phosphorylated VEGFR2 (pR2) [23] and pERK [22] responses induced by VEGF were obtained using human umbilical vein endothelial cells (HUVECs). The FGFR1 and HSGAG levels on various cell types are fairly consistent; specifically, approximately 10^4^ to 10^5^ molecules/cell for Balb/c3T3 [24], osteoblasts and bone marrow stromal cells [25], bovine aortic endothelial cells (BAECs) [26], and NCI-H1730 [21]. Thus, we assume the FGFR1 signaling pathway for NCI-H1730 is the same as in endothelial cells.

**Figure 1.**
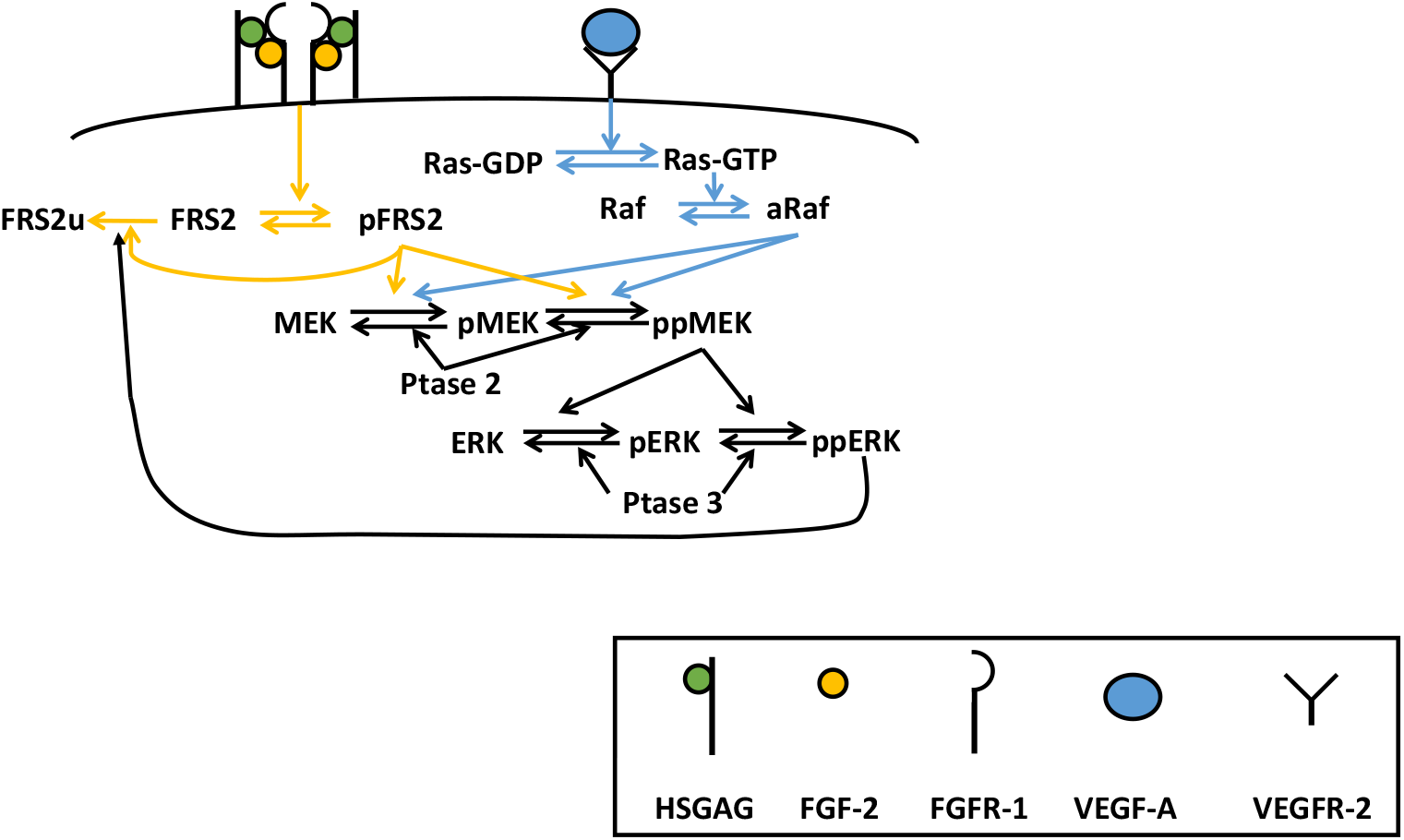
Schematic of FGF and VEGF signaling network. Signaling is induced by the growth factors binding to their receptors, culminating with phosphorylation of ERK, through the MAPK cascade. MAPK signaling is initiated through the activation of Raf and FRS2 by VEGF and FGF, respectively. The FGF:HSGAG:FGFR1 complex dimerizes and leads to phosphorylation of FRS2 (pFRS2). VEGF binds VEGFR2 to activate Ras, forming Ras-GTP, which further activates Raf (aRaf). Both aRaf and pFRS2 are able to phosphorylate MEK at two sites, and doubly phosphorylated MEK (ppMEK) further phosphorylates ERK. ppERK provides negative feedback on the FGF pathway, as it promotes ubiquination of FRS2 (FRS2u).

The fitted model shows a good match to the training data (Figure 2). It can capture the biphasic pERK response caused by FGF stimulation, which has been reported by Kanodia *et al*. (Figure 2 A, C): pERK increases as the FGF concentration increases from low to intermediate levels, and decreases with increasing FGF at high concentrations. The decrease in pERK is caused by the competitive binding of FGF to HSGAG and FGFR [21]. Our fitted model recapitulates this biphasic response at the simulated time points. Also, VEGF-induced upstream (pVEGFR2) and downstream (pERK) dynamics have good agreement with experimental measurements (Figure 2B, D) [22, 23]. For the best 16 fits, the weighted errors range from 17.4 to 18.6 (Table S1). In addition, the fitted parameters have good consistency, as the estimated values for each parameter are all within two orders of magnitude, and many parameters have an even tighter range (Figure S1).

**Figure 2.**
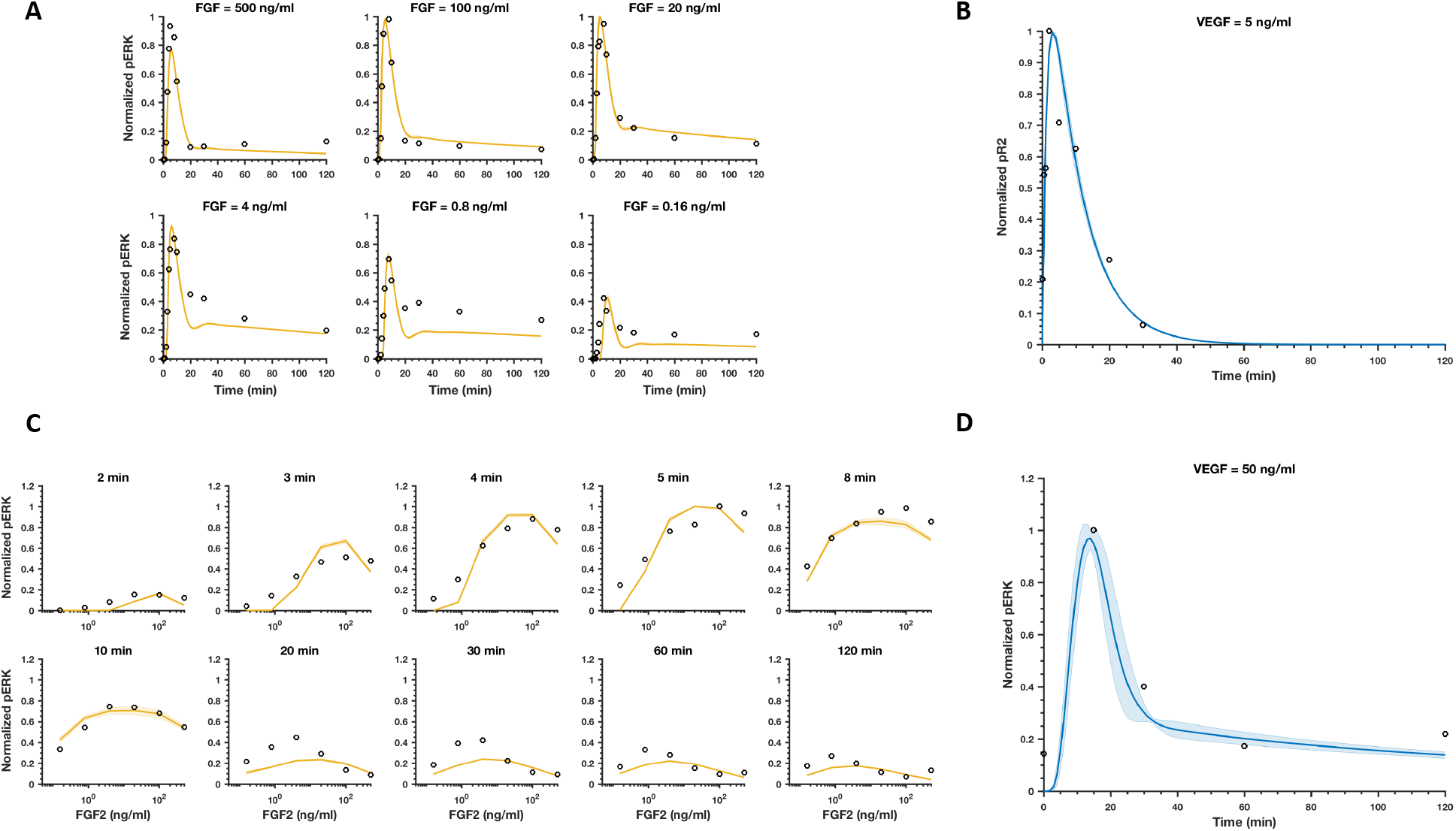
Model comparison to training data for FGF or VEGF stimulation. (A) Normalized pERK dynamics in response to FGF concentrations ranging from 0.16 to 500 ng/ml. (B) Normalized VEGFR2 phosphorylation time course following stimulation with 5 ng/ml VEGF. (C) Dose response of pERK for FGF stimulation. (D) Normalized ERK phosphorylation time course upon stimulation with 50 ng/ml VEGF. The circles are experimental data. Curves are the mean values of the 16 best fits. Shaded regions show standard deviation of the fits.

### Model validation shows good qualitative and quantitative agreement with experimental results

To validate the model, we compared the model predictions to additional experimental data. We first applied heparin perturbation to the trained model to reproduce another set of data by Kanodia *et al*. Heparin is a soluble source of heparin sulfate glycosaminoglycans (HSGAGs), and it competes with HSGAGs to bind with FGF, interfering with FGF-induced signaling (Figure S2) [21]. It has been reported that additional heparin increases FGF-induced ERK phosphorylation at high FGF concentrations, and decreases FGF-induced ERK phosphorylation at low FGF concentrations in two hours [21]. We validated the model by expanding it to include heparin binding and comparing to the experimental measurements. We added 500 µg/ml of heparin, the same concentration used in experiments [21], and found that the difference of predicted pERK responses upon FGF stimulation with and without heparin within two hours exhibits the trend observed to occur experimentally: at high FGF concentrations, the difference between in the presence and absence of heparin is greater than zero, while the difference is less than zero at lower FGF concentrations (Figure 3A). This shows a qualitative agreement with experimental observations.

A separate set of experimental measurements of phosphorylated ERK by the stimulation of 10 ng/ml FGF conducted using BAECs [27] was extracted to further validate FGF-induced endothelial signaling. Our model also has a good agreement with this experimental data (Figure 3B), which further confirms that this model can be used to predict endothelial cell signaling.

**Figure 3.**
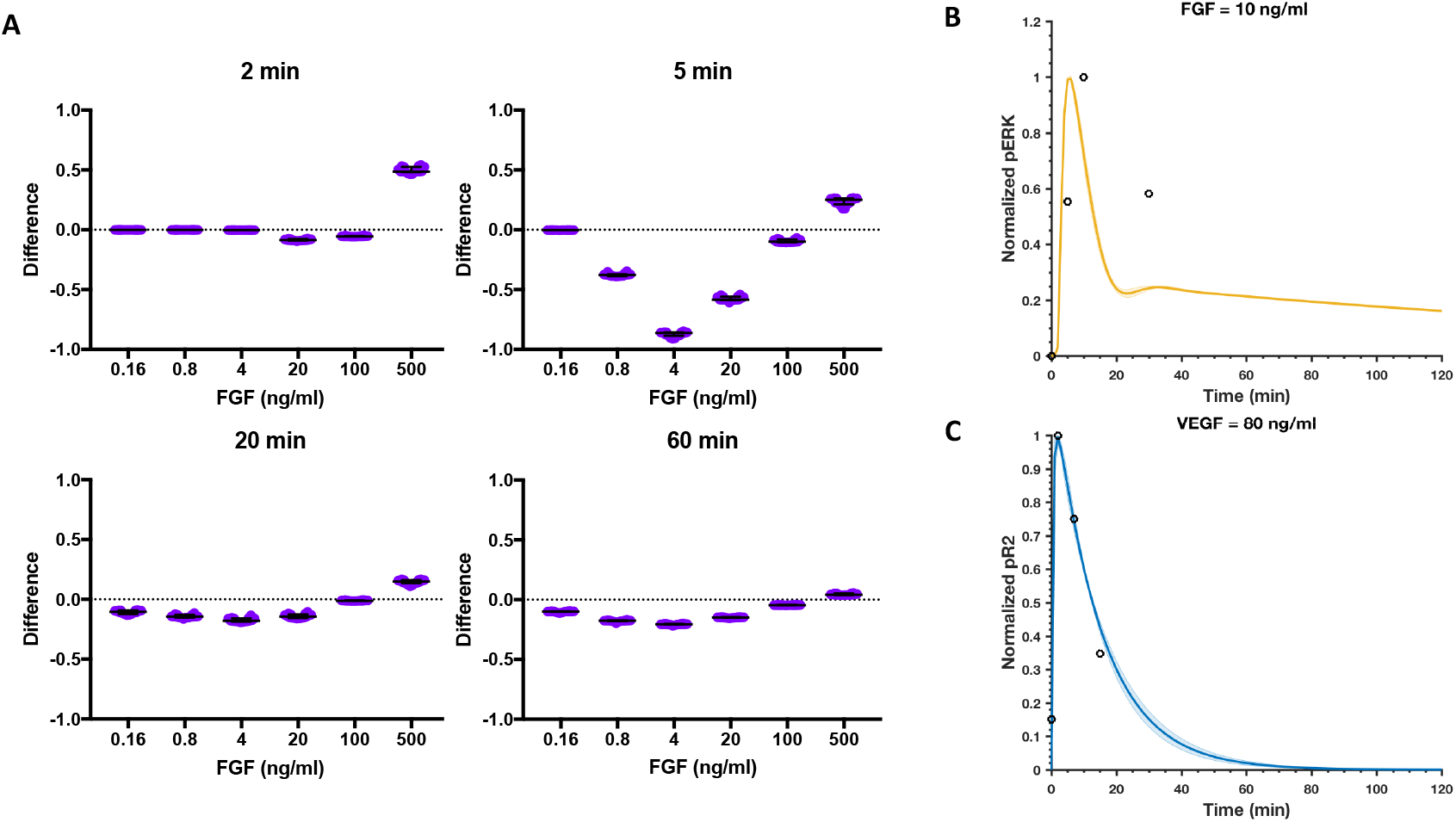
Model comparison to validation data. (A) The differences in pERK induced by stimulation of FGF with and without 500 µg/ml heparin at four time points are predicted. (B) Normalized pERK by the stimulation of 10 ng/ml FGF. Circles are BAEC experimental data. (C) Normalized VEGFR2 phosphorylation time course upon stimulation with 80 ng/ml VEGF. Circles are HUVEC experimental data. Curves are the mean values of the 16 best fits from the model training. Shaded regions show standard deviation of the fits.

We also extracted an independent set of experimental measurements of the phosphorylated VEGFR2 response by the stimulation with 80 ng/ml VEGF in HUVECs [28] to validate the VEGF-induced signaling. Our model quantitatively matches the experimental data (Figure 3C). Overall, we find that our trained model can generate reliable predictions for the endothelial intracellular signaling response stimulated by FGF or VEGF. Thus, we used the best fits (based on model training and validation) in subsequent predictions and analyses.

### FGF produces a greater angiogenic response than VEGF when considering the maximum ERK phosphorylation produced

Using the validated model, we first studied the effects of FGF and VEGF individually on pERK. The model predicts that FGF is more potent in promoting ERK phosphorylation, compared to VEGF, at equimolar concentrations. When FGF concentration is varied from 0.01 nM to 2 nM, the maximum pERK ranges from 4×10^5^ molecules/cell to 8×10^5^ molecules/cell, while VEGF induces a maximum of 2×10^-2^ molecules/cell to 1×10^5^ molecules/cell pERK for the same concentration range (Figure 4A). For example, on average (across the 16 best fits), the maximum pERK induced by 0.5 nM FGF is 8×10^5^ molecules/cell, while 0.5 nM VEGF induces a maximum pERK of 9×10^2^ molecules/cell. Thus, FGF produces a maximum ERK phosphorylation that is approximately three orders of magnitude higher than that induced by VEGF. Furthermore, the maximum pERK is more sensitive to increasing the VEGF concentration, as compared to FGF. The maximum pERK increases steadily with increasing VEGF stimulation, while maximum pERK remains relatively constant as the level of FGF stimulation increases. For the FGF pathway, the phosphorylated trimeric complex of FGF, FGFR, and HSGAG binds to and phosphorylates FRS2, and phosphorylated FRS2 (pFRS2) leads to the phosphorylation of MEK. The resulting doubly phosphorylated MEK (ppMEK) further mediates the phosphorylation of ERK (Figure 1). We found that even with 0.01 nM FGF stimulation, FRS2 is rapidly depleted (Figure S3A). Therefore, FRS2 level limits the FGF-induced ERK phosphorylation. Increasing FGF concentration 200-fold (from 0.01 nM to 2 nM) only doubles the maximum pERK (from 4×10^5^ molecules/cell to 8×10^5^ molecules/cell), again due to the shortage of FRS2. On the other hand, for the VEGF pathway, phosphorylated VEGFR2 produces Ras-GTP, which activates Raf. The activated Raf (aRaf) mediates phosphorylation of MEK. As in the FGFR pathway, ppMEK mediates ERK phosphorylation (Figure 1). The model predicts that there is enough Raf and MEK supply even upon stimulation with a high concentration of VEGF (2 nM), as shown in Figure S3B. Thus, the maximum pERK increases significantly with increasing VEGF concentration, compared with FGF-induced ERK phosphorylation. Furthermore, the maximum pERK induced by FGF or VEGF gets closer as VEGF concentration increases (Figure 4A).

**Figure 4.**
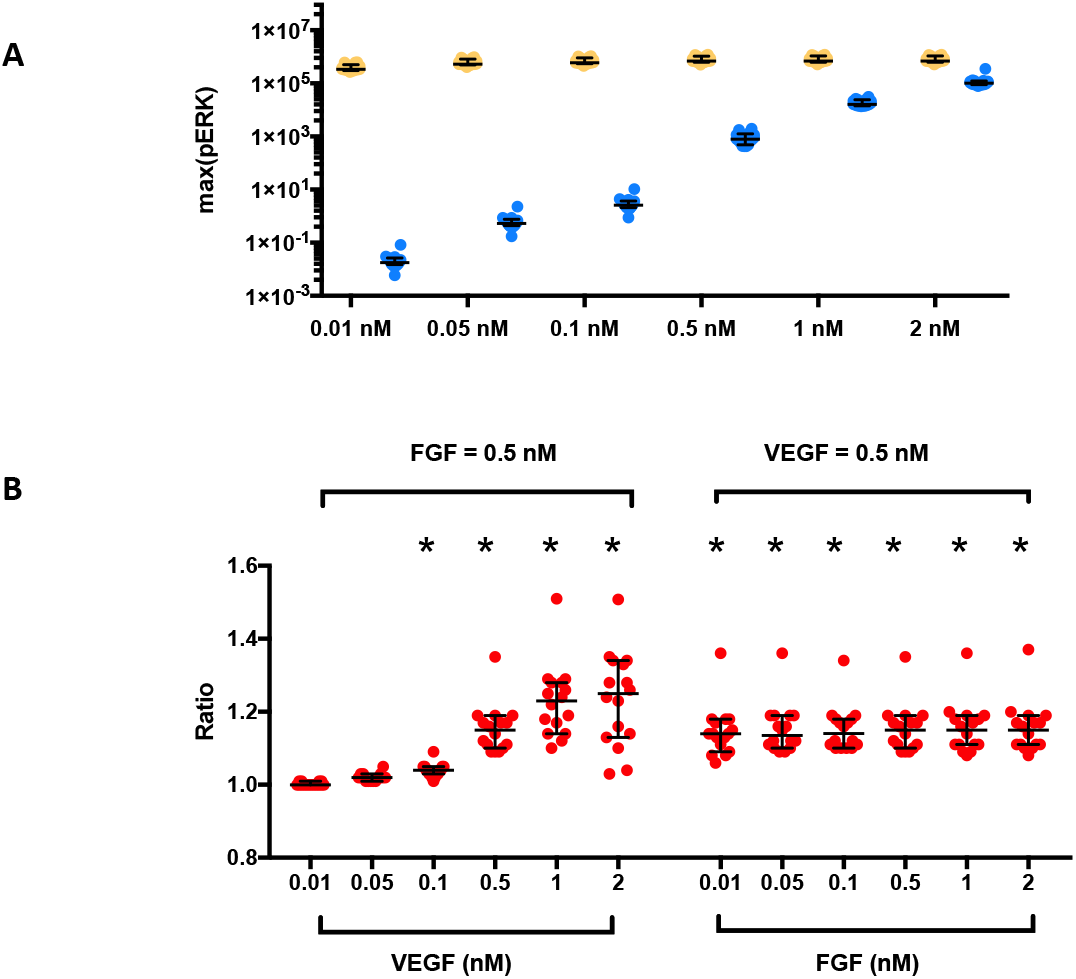
Predicted maximum pERK response. (A) Maximum pERK in response to FGF (yellow) or VEGF (blue) concentrations varying from 0.01 nM to 2 nM. (B) Ratio, *R*, of combination effects to the summation of individual effects in response to FGF and VEGF. Each dot represents one fit. Asterisk indicates statistically significant difference compared with one (*p*<0.05). Bars are median ± 95% confidence interval.

The model predicts that the main reasons why FGF induces a greater maximum pERK response compared to VEGF are related to differences at the receptor level. Despite depletion of FRS2, high FGFR levels enable robust FGF-mediated signaling. FGFR density is much higher than VEGFR2 density (20,000 molecules/cell compared to 1,000 molecules/cell). Additionally, the trafficking parameters (internalization, recycling, and degradation rates) values for FGFR are lower than the corresponding VEGFR2 trafficking parameters (Figure S4). Although internalized VEGFR2 molecules are recycled back to the surface more rapidly than FGFR molecules, VEGFR2 is internalized and degraded more rapidly than FGFR. Additionally, the dynamics of FGF receptors and VEGFR2 in their signaling, internalized and degraded forms upon stimulation with 0.5 nM FGF or 0.5 nM VEGF (Figure S5) indicate that more FGFR is available to signal instead of being internalized or degraded (non-signaling), compared to VEGFR2.

### The combination of FGF and VEGF has greater effects in inducing maximum pERK than the summation of the individual effects

We next studied the combination effects of FGF and VEGF in inducing maximum pERK. Here, we define a ratio comparing the combination effects to the individual effects. Specifically, this ratio is the maximum pERK obtained with co-stimulation of FGF and VEGF over the summation of the maximum pERK for FGF and VEGF stimulation individually (see Methods for more details). In Figure 4B (left panel), 0.5 nM FGF in combination with intermediate to high VEGF concentrations (0.1–2 nM) can produce a significantly greater maximal pERK response than the summation of their individual effects, as indicated by the ratios being significantly greater than one (*p*<0.05). The ratios for combinations of VEGF concentrations at 0.01 or 0.05 nM with 0.5 nM FGF are slightly greater than one; however those differences are not statistically significant. Stimulation with 0.5 nM VEGF in combination with a FGF concentration as low as 0.01 nM can exhibit greater combined effects than the summation of their individual effects. For these cases, the ratios are all significantly greater than one (Figure 4B, right panel).

The effects of FGF and VEGF co-stimulation are more sensitive to VEGF, as compared to FGF. That is, increasing the VEGF concentration increases the ratio, while the ratio does not change with varying FGF concentrations. Additionally, the combination of FGF with VEGF shows a more additive response at low VEGF concentrations (< 0.1 nM). At higher concentrations (> 0.1 nM), increasing VEGF concentration increases the ratio (Figure 4B).

Overall, the model predictions show that combinations of FGF and VEGF produce more ERK phosphorylation, compared to their individual effects. Additionally, the model indicates that VEGF-induced maximum pERK is more sensitive to varying the ligand concentration than FGF-induced maximum pERK, both for stimulation with VEGF or FGF alone and for co-stimulation.

### The combination of FGF and VEGF shows a fast and sustained pERK response

*The combination of FGF and VEGF exhibits a fast pERK response*. In addition to studying the magnitude of the predicted pERK level upon FGF and VEGF mono-and co-stimulation, we investigated the timescale of the pERK response. First, we analyzed how fast ERK is activated in response to the stimulation of FGF or VEGF individually, and we found that FGF generally produces a faster response than VEGF stimulation at the same concentrations. Here, we characterize the timescale of the response in terms of the time it takes to reach maximum pERK, termed “T1” (see Methods for more details). At lower concentrations (0.01 – 0.5 nM), the T1 values for FGF and VEGF are not significantly different. However, at higher concentrations (0.5 – 2 nM), FGF shows a significantly faster T1 response than VEGF. Specifically, for FGF concentrations ranging from 0.5 to 2 nM, the induced pERK response peaks within six minutes, while for the same range of VEGF concentrations, pERK reaches its peak value within 8 to 22 minutes (Figure 5A). Together with Figure 4A, the model predicts that FGF can induce a greater amount of ERK phosphorylation within a shorter period of time, compared to VEGF.

As for the combination effects, we found that when the VEGF concentration is varied from 0.01 nM to 0.5 nM, co-stimulation with 0.5 nM FGF significantly speeds up ERK phosphorylation, compared to VEGF stimulation alone (Figure 5A). For 0.5 nM VEGF stimulation, increasing the FGF concentration decreases T1, compared to VEGF stimulation alone. This decrease in T1 is significantly different than VEGF stimulation alone for FGF concentrations greater than 0.5 nM (Figure 5A).

**Figure 5.**
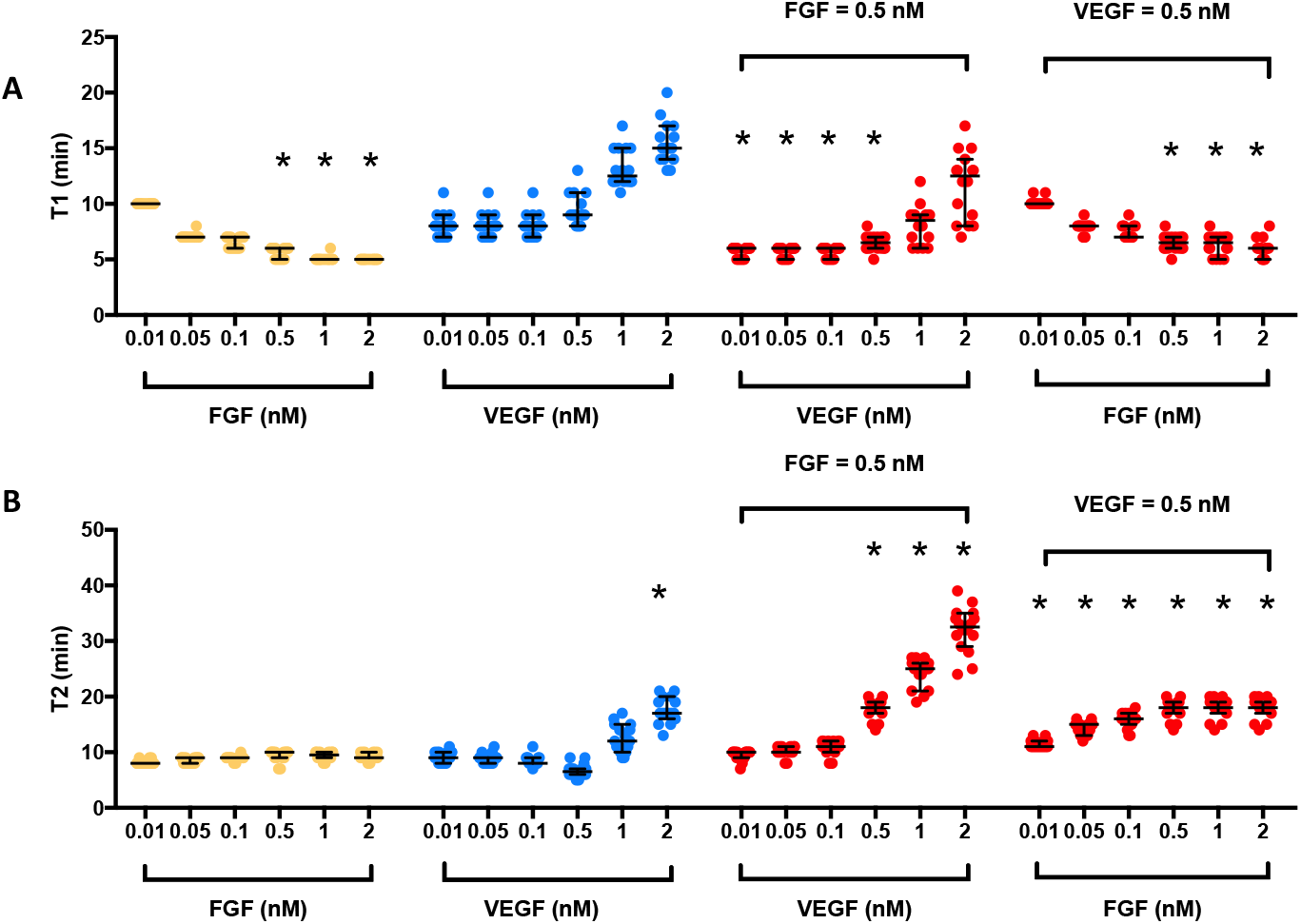
Predicted time response of pERK following stimulation by FGF, VEGF, and their combination. (A) *T1*, time to reach the maximum pERK in response to growth factor stimulation. Asterisk indicates statistically significant difference compared to corresponding VEGF concentration (*p*<0.05). (B) *T2*, time for pERK maintain above half maximum in response to treatments. Each dot represents one fit. Asterisk indicates statistically significant difference compared to corresponding FGF concentration (*p*<0.05). Yellow: FGF; Blue: VEGF; Red: combination. Bars are median ± 95% confidence interval.

Overall, these results indicate that pERK responds faster with FGF stimulation, as compared to VEGF stimulation. Again, this is largely due to more FGFR being available to signal compared to VEGFR2 (Figure S5).

*The combination of FGF and VEGF induces sustained pERK response*. We explored how long ERK can remain phosphorylated above its half-maximal value, termed “T2” (see Methods for more details), as another means of characterizing the timescale of the ERK response. The values of T2 for FGF and VEGF stimulation alone with concentrations ranging from 0.01 nM to 1 nM are not significantly different (T2 is approximately 9 minutes). However, a higher VEGF concentration produces a more sustained pERK response. Specifically, 2 nM VEGF shows significantly higher T2 (18 minutes on average) than 2 nM FGF stimulation (9 minutes on average) (Figure 5B).

Regarding the combination effects, we found that with 0.5 nM FGF, VEGF concentrations greater than 0.5 nM are able to maintain ERK phosphorylation above its half-maximal value significantly longer, compared to FGF stimulation alone at the same concentrations. That is, T2 is significantly greater for combinations of 0.5 nM VEGF with FGF concentrations ranging from 0.01 nM to 2 nM, compared to FGF or VEGF stimulation alone (Figure 5B).

To identify the reasons why pERK shows a more transient dynamic in response to FGF stimulation compared to VEGF stimulation at certain concentrations (>0.5 nM), we compared the levels of the intermediate signaling species following stimulation of FGF or VEGF alone. As shown in Figure S3A, there is a shortage of FRS2 that limits the phosphorylation of FRS2 stimulated by FGF and ultimately leads to a transient pERK response. In contrast, Figure S3B shows enough Raf and MEK supply for VEGF stimulation at higher concentrations. However, as the VEGF concentration increases and more ppMEK is produced, more phosphatase (Ptase2) is utilized. Thus, at higher VEGF concentrations, there is not enough available Ptase2 to dephosphorylate pMEK and ppMEK (Figure S3B). This leads to more sustained ppMEK dynamics and subsequently, a more sustained ppERK response.

For the co-stimulation of FGF and VEGF, the depletion of FRS2 still limits production of ppMEK with FGF stimulation (Figure S6). However, signaling through VEGFR compensates for this limitation (Figure S6). Specifically, the sustained pERK response is possible since the ppMEK response is the net result of both FGF and VEGF stimulation.

### FGF plays a dominant role in the combination effect of FGF and VEGF in inducing maximum pERK and time to reach maximum pERK

To further understand the interactions of FGF and VEGF mechanistically, we examined relevant species’ dynamics and reaction rates in detail. The time course for ppERK shows that the average maximum ppERK levels in response to 0.5 nM FGF, 0.5 nM VEGF, and their combination are 1.1×10^4^, 2.5×10^-3^, and 1.5×10^4^ molecules/cell, respectively (Figure 6A). These predictions demonstrate the results presented above, showing that FGF induces a higher maximum pERK than VEGF, and that the combination effects are greater than individual effects. Additionally, the pERK response to FGF and VEGF co-stimulation shows features of individual FGF and VEGF stimulation: faster and more sustained response, compared to stimulation with VEGF or FGF alone (Figure 5).

**Figure 6.**
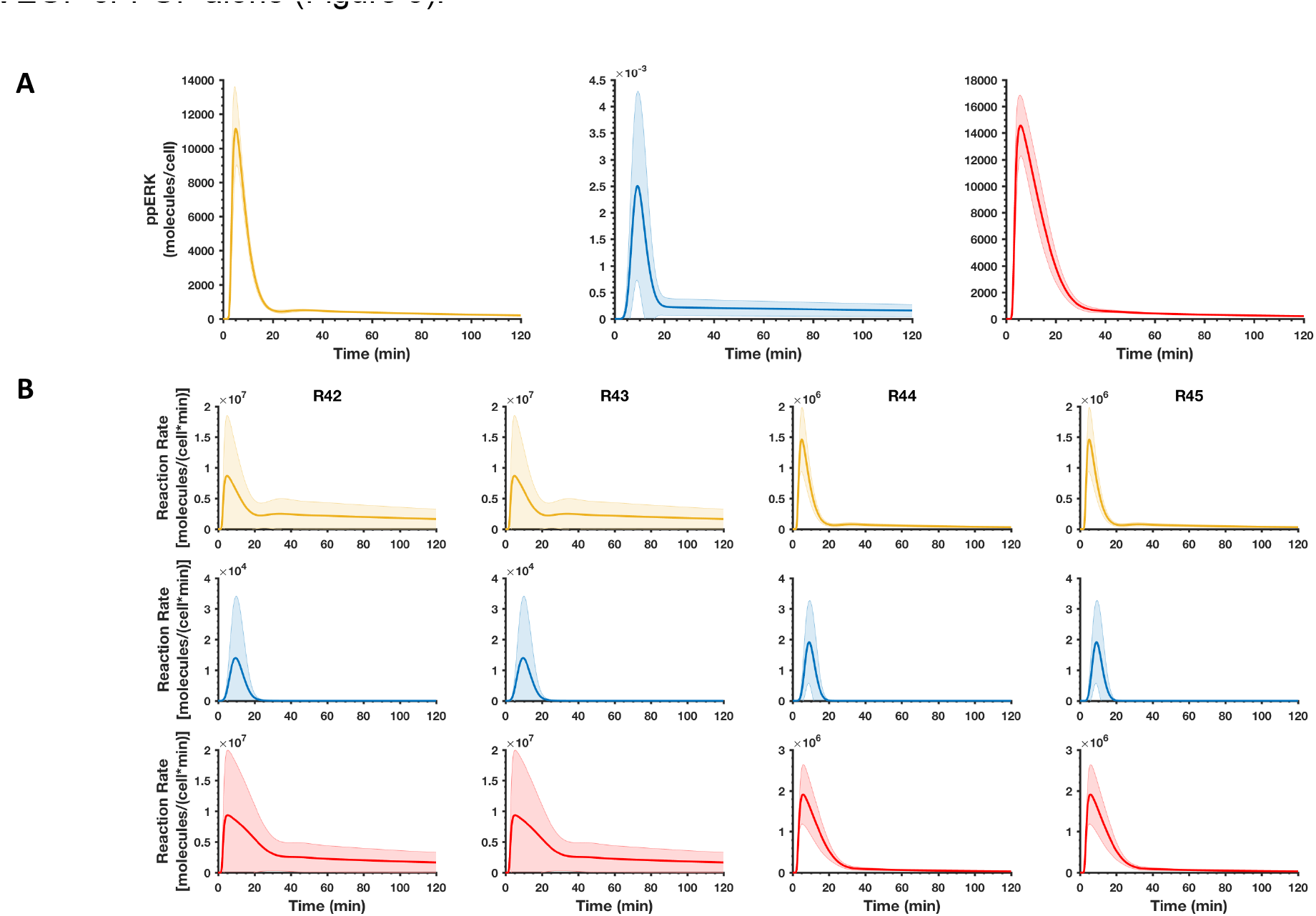
Reaction rates in response to 0.5 nM FGF, 0.5 nM VEGF, and their combination stimulation. (A) Time course for ppERK dynamics. (B) Reaction rates for ERK and pERK phosphorylation. R42: ERK + ppMEK ↔ ERK:ppMEK; R43: ERK:ppMEK pERK:ppMEK → ppERK+ppMEK. Yellow: FGF; Blue: VEGF; Red: combination. Curves are the mean values of the 16 best fits, and shaded regions show standard deviation of the fits.

To explore the individual contributions to the combination effects, we investigated the reaction rates for MEK and ERK phosphorylation induced by FGF and VEGF. In Figure S7, FGF is dominant in phosphorylating MEK and pMEK for FGF and VEGF co-stimulation. Specifically, the reaction rates for MEK phosphorylation with 0.5 nM FGF stimulation are more than one order of magnitude higher than for 0.5 nM VEGF stimulation (Figure S7A, B). This leads to the formation of more ppMEK, the species that mediates ERK phosphorylation. Thus, these increased reaction rates significantly affect the levels and dynamics of pERK. For combinations of FGF and VEGF, FGF is dominant in phosphorylating MEK and pMEK, as R25 through R28 have much higher reaction rates than R34 through R37 (Figure S7C). Thus, the time to reach maximum pERK is more dependent on FGF-mediated stimulation. We note that the rates of R34 and R35 for FGF and VEGF co-stimulation are slightly lower than VEGF stimulation alone. This is due to the competition for MEK between the FGF and VEGF pathways, where more MEK is bound by FGF-induced pFRS2 than aRaf from the stimulation of VEGF. However, the rates of R36 and R37 are much higher for combinations of FGF and VEGF, compared to VEGF stimulation alone. This is because more pMEK, the substrate for reactions R36 and R37, is produced by the co-stimulation.

Furthermore, we studied the reactions directly leading to ERK phosphorylation (Figure 6B). The reaction rates of R42 and R43 are almost identical during the first 10 minutes for FGF alone compared to the combination of FGF and VEGF. However, later (10 to 60 minutes), the reaction rates for combined stimulation of FGF with VEGF exceed the rates for FGF alone. Additionally, the reaction rates remain at a slightly higher level, leading to a more sustained response. The rates of the VEGF-stimulated ERK phosphorylation reactions are much lower in comparison with FGF or the combination of FGF and VEGF, for all simulated times.

These model predictions for the estimated reaction rates hold true even when VEGF concentration is increased to 2 nM (data not shown). Thus, the effect of FGF is predicted to dominate the pERK response for combined stimulation of FGF and VEGF, based on several characteristics of pERK dynamics.

### Increasing VEGFR2 density can compensate for the relatively low ERK phosphorylation induced by VEGF

At its baseline level, the density of VEGFR2 is 20 times lower than FGFR1. This large difference contributes to the predicted results presented above. Experimental measurements of receptor expression show that some subpopulations of tumor endothelial cells (ECs) have high receptor levels: 13% of tumor-derived ECs have 7,500 VEGFR2 molecules/cell after three weeks of tumor growth, and 5% of the tumor-derived ECs have 16,200 VEGFR2 molecules/cell after six weeks of tumor growth [29]. Therefore, we sought to understand the effects of varying the VEGF receptor density on the predicted pERK response.

We found that the maximum pERK induced by VEGF increases when VEGFR2 density increases (Figure 7A). At equimolar concentrations of FGF and VEGF stimulation (0.5 nM), increasing VEGFR2 density by five-fold can increase the maximum pERK level to the same order of magnitude as FGF stimulation alone (Figure 7B). Specifically, the model predicts that maximum pERK induced by the combination of FGF and VEGF increases more than 90% when VEGFR2 is increased by five-fold. In contrast, decreasing VEGFR2 by ten-fold leads to an 11.5% decrease in maximum pERK induced by the combination of FGF and VEGF. Thus, VEGFR2 density significantly impacts ERK phosphorylation with FGF and VEGF co-stimulation. In addition, the maximum ppERK level is higher upon stimulation by VEGF, compared to FGF, when VEGFR2 density is increased (Figure S8A). Moreover, although the reaction rates for ERK phosphorylation (R42 and R43) by stimulation of FGF are slightly higher than VEGF during the first 10 minutes, VEGF induces higher rates between 10 to 60 minutes (Figure S8B). In addition, VEGF exhibits faster phosphorylation for pERK (R44 and R45) than FGF (Figure S8B).

This indicates that the effect of VEGF is dominant in the combination effect when VEGFR2 density is increased by five-fold.

In Figure 7C, as VEGFR2 density decreases, the ratio characterizing the ERK signaling response with a combination of FGF and VEGF compared to the summation of their individual effects becomes closer to one. This suggests that the combination effect is nearly additive. This is because reducing VEGFR2 makes the effect of VEGF stimulation negligible. Thus, the ratio is approximately one. The ratio increases when VEGFR2 density is increased by two-fold, which indicates a stronger combination effect. However, the summation of individual effects surpasses the combination effects (the ratio is less than one) when VEGFR2 density is more than five-fold higher than the baseline level. The reason for this is due to the competition between FGF and VEGF for downstream resources. Since the increase of VEGFR2 impacts downstream signaling, when VEGFR2 is increased by ten-fold, ERK is depleted (Figure S9B). This makes the combination effects less than the individual effects, making the ratio less than one. Finally, by increasing VEGFR2, VEGF more strongly impacts the dynamics of pERK, as both T1 (Figure 7D) and T2 (Figure 7E) increase with increasing VEGFR2 density.

### ERK phosphorylation induced by VEGF can be promoted by decreasing VEGFR2 internalization and degradation rates

Because FGFR trafficking parameter values are lower than the corresponding VEGFR2 trafficking parameters (Figure S4), we explored the role of trafficking parameters in pERK response. Specifically, we investigated the effects of decreasing the VEGFR2 trafficking parameters individually or together to the same level as FGFR trafficking parameters (Figure 8).

We found that decreasing the internalization rates of the free and bound forms of VEGFR2, individually or in combination, with 0.5 nM VEGF leads to a significant increase in maximum ERK phosphorylation (Figure 8A, “*fitted*” compared to “*k_intf*”, “*k_intb*”, or “*k_int*”). In fact, by decreasing VEGFR2 internalization (“*k_int*”) to be the same as the FGFR internalization rate, the maximal VEGF-mediated ERK phosphorylation reaches the same magnitude as the response induced by 0.5 nM FGF. This qualitative trend is expected, since decreasing receptor internalization rates makes more VEGFR2 available on the cell surface to bind to the ligand, inducing downstream signaling. However, the model provides detail about the quantitative effects of these changes.

Decreasing the recycling rates of the free and bound forms of VEGFR2 individually or in combination significantly decreases ERK phosphorylation for 0.5 nM VEGF stimulation (Figure 8A, “*fitted*” compared to “*k_recf*”, “*k_recb*”, or “*k_rec*”). A lower receptor recycling rate makes more non-signaling internalized VEGFR2 remain inside the cell longer and recycles available VEGFR2 to the cell surface more slowly. Together, these effects limit ERK phosphorylation. Interestingly, changing the recycling rate leads to a wider range of responses compared to changing internalization or degradation rates, especially for recycling of free VEGFR2 (Figure 8A).

Lowering the rate at which the free form of VEGFR2 is degraded significantly increases the maximum ERK phosphorylation induced by 0.5 nM VEGF to the same magnitude of the maximal pERK level induced by 0.5 nM FGF alone (Figure 8A, “*fitted*” compared to “*k_degf*”, “*k_degb*”, or “*k_deg*”). Lower free receptor degradation rate makes more VEGFR2 available to promote signaling.

We also examined the timescales of the ERK response. The model predicts that decreasing the trafficking rates slows down the dynamics of VEGF-induced ERK phosphorylation (Figure 8B, C), both in terms of T1 and T2. Furthermore, we studied the effects of changing VEGFR2 trafficking parameters on the combination effects. We found that the combination of 0.5 nM FGF and 0.5 nM VEGF with lower VEGFR2 trafficking parameters has similar results as 0.5 nM VEGF-induced pERK. That is, decreasing VEGFR2 internalization and degradation rates leads to greater ERK phosphorylation (Figure S10). Overall, lower VEGFR2 trafficking parameters leads to an increased impact of VEGF in the combination effects.

## Discussion

We developed an intracellular signaling model of the crosstalk between two proangiogenic factors, FGF and VEGF. The molecular-detailed model represents the reaction network of interactions on a molecular level, based on reactions documented in literature. The kinetic parameters are taken from experimental measurements, where available. Unknown parameters were estimated by fitting the model to experimental data. Additionally, we validated the model using three separate sets of data.

This is a novel model of FGF and VEGF interactions, taking into account previous modeling work [11, 21], with a focus on the MAPK cascade and pERK response. The fitted model predicts the pERK response upon stimulation by FGF and VEGF, alone and in combination. We particularly focus on the pERK response since pERK promotes cell proliferation [19], one aspect of early-stage of angiogenesis. Additionally, it has been shown that pERK is mostly found in active rather than quiescent endothelial cells [20]. Thus, in this study we use pERK response as an indicator for pro-angiogenic signaling. Overall, FGF is predicted to potently and rapidly promote ERK phosphorylation compared to VEGF stimulation. Altogether, the model shows that the pERK level in response to FGF, VEGF, and their combination is dose-dependent and that some combinations induce greater maximum ERK phosphorylation than the summation of their individual effects.

The model predictions are consistent with several experimental studies. Multiple experiments show that FGF induces the same level of angiogenic response at lower concentration in comparison to VEGF [30-32], and their combination induces greater angiogenic responses [13]. The model also predicts that the density of VEGFR2 and the rates of internalization, degradation, and recycling of VEGFR2 contribute to the lower ERK phosphorylation induced by VEGF. Specifically, the model predicts that increasing VEGFR2 concentration on the cell surface can compensate for low VEGF-induced ERK phosphorylation. This is particularly relevant, as endothelial VEGF receptors expression is upregulated in tumors [33], and can reach nearly 2×10^4^ molecules/cell in tumor-derived endothelial cells after six weeks of tumor growth [29]. Additionally, the model predicts that decreasing VEGFR2 internalization and degradation rates can increase the impact of VEGF in combination effects. This result complements experiments showing that receptor trafficking plays a critical role in angiogenic signaling [34]. Overall, our molecular-detailed model helps synthesize these experimental data and observations related to VEGF-and FGF-stimulated signaling.

One application of our work is that the model can also be linked with computational models that predict events on the cellular scale. Our model culminates with ERK activation, complementing published models that substantially simplify the intracellular signaling and focus on specific cellular behavior, such as proliferation [35], the probability of sprout formation and the speed of vessel growth [36], or tumor growth [37]. However, these models reduced the intracellular signaling network such that the output signal is simply linearly proportional to the fraction of bound receptors. In comparison, our mechanistic model considers intracellular signaling and quantitatively analyzes pERK response, which could be a better indicator for these cellular behaviors. For example, Hendrata and Sudiono constructed a computational model that includes molecular, cellular, and extracellular scales to study tumor apoptosis [37]. Our model can be utilized in combination with such models to more accurately predict cellular behavior.

Our model can also aid in studying the efficiency of pro-or anti-angiogenic therapies. Some pro-or anti-angiogenic treatments have not been very effective, particularly those targeting only a single signaling family [7, 8]. However, targeting both FGF and VEGF may be a promising strategy, given the potential synergistic effects predicted by our model and demonstrated in experimental studies. In fact, multiple groups have reported interesting interactions between FGF and VEGF [12, 13, 38, 39]. This crosstalk may be exploited to aid in angiogenesis-based therapies, and our model can be helpful in understanding their interactions and combination effects. Model predictions for species’ dynamics and reaction rates provide mechanistic insight into FGF and VEGF interactions. Our predictions show that the low success in targeting VEGF alone could be due to low receptor numbers and fast internalization, recycling and degradation. Although FGF has greater effects in inducing ERK phosphorylation, its effects can be enhanced by the addition of VEGF. Thus, our model can be used to investigate the efficiency of targeting both FGF and VEGF as an alternative strategy.

Our model is the first to combine the signaling networks of FGF and VEGF, providing novel quantitative insight into the effect of combined FGF and VEGF treatment. However, we recognize some limitations in our model. Firstly, this model does not include FGF activated Ras-Raf signaling because the protein-protein interactions in this pathway are still not clear. We did implement alternative network structures with greater detail from FGF to ERK and refit the data; however, these models did not provide a better fit. Thus, we retained the model shown in Figure 1. Second, we assumed that the internalized and degraded phosphorylated receptors would not signal. However, data suggest that VEGFR2 may signal even when internalized [40], which can be considered in future studies. Additionally, all bound forms of FGFR1 are assumed to have the same internalization, recycling, and degradation rates as a simplification and because there are conflicting values reported in literature [24, 41]. We tried various FGFR1 trafficking parameters; however, this did not significantly change the model predictions or our overall conclusions. Third, this model only includes VEGFR2, although VEGF binds to VEGFR1 and neuropilin-1 (NRP1). These receptors also contribute to angiogenesis and may be incorporated into the model in future studies. Finally, we studied pERK response in two hours. We omit ligand secretion and protein degradation during this time and do not predict long-term responses. In the future, we can expand our model to predict the cellular response over a longer period of time.

## Conclusions

In summary, our molecular-detailed model quantifies ERK phosphorylation upon stimulation by two major pro-angiogenic factors, FGF and VEGF, and provides insights into the molecular interactions between these proteins. Specifically, the model predicts the combination effects of FGF and VEGF on ERK phosphorylation and quantitatively shows the magnitude and time scale of the pERK response. Because of the complexity of this biological system, it may be challenging to get a comprehensive understanding of system using experiments that only focus on a few molecular species. Our computational modeling provides a quantitative framework to explore the system as a whole, generating novel mechanistic insight and complementing experimental studies.

## Methods

### Model construction

We constructed a molecular-detailed biochemical reaction network including FGF, VEGF, and their receptors FGFR1 and VEGFR2 (Figure 1). Signaling is induced by the growth factors binding to their receptors, culminating with phosphorylation of ERK, through the MAPK cascade. MAPK signaling is initiated through the activation of Raf and FRS2 by VEGF and FGF, respectively. Activated Raf (aRaf) and phosphorylated FRS2 (pFRS2) phosphorylate MEK at two sites, and doubly phosphorylated MEK (ppMEK) further phosphorylates ERK. The molecular interactions involved in the network are illustrated in Figure 1. The model is a novel advancement of published computational models [11, 21]. Specifically, we adapted the competition of FGF and HSGAG to the binding of FGFR1 and the feedback loop from pERK to FRS2 from the model by Kanodia *et al*., and expanded the model by including FGFR trafficking (internalization, recycling, and degradation) and accounting for both singly-and doubly-phosphorylated MEK and ERK into consideration. On the other hand, we simplified the model of VEGF-induced ERK phosphorylation pathways from Tan and coworkers; specifically, we only include Ras activation either from Shc-independent or Shc-dependent pathways [11]. Thus, we improved upon the previous works to capture the major steps of FGF-and VEGF-induced ERK phosphorylation and better understand their interactions.

The network is implemented as an ordinary differential equation (ODE) model using MATLAB. The main model includes 70 reactions, 72 species, and 75 parameters. The initial conditions and parameter values are listed in Tables S2-S4. All reactions are assumed to follow the law of mass action. Receptor internalization, recycling, and degradation are considered in the model. Because the simulated time is within two hours, we do not consider the degradation of the ligands or signaling species. The final model is available in Additional File 3. We also implement a modified model that includes heparin to validate the estimated model parameters (described below).

### Data extraction

Data from published experimental studies [21-23,27, 28] were used for parameter fitting and model validation. The Western blot data was extracted using ImageJ. Experimental data from plots was extracted using function *grabit*.

### Model parameters

The trafficking parameters for VEGFR2 and the parameters and initial values that are involved in the overlap of FGF and VEGF signaling pathways were estimated by fitting the model 72 times to experimental data using Particle Swarm Optimization (PSO). A total of 39 parameters and initial values were estimated in the fitting (Table S1, and also highlighted in red in Tables S3-S4). All other parameters were taken from published literature [11, 21, 24]. The parameters characterizing the overlapping MAPK pathway were chosen for fitting because while FGF and VEGF upstream parameters are well documented individually in literature, a uniform set of parameters for their interactions is needed for this combined model. The VEGFR2 trafficking parameters were fitted as they have been shown to significantly affect ERK activation [11].

PSO starts with a population of initial particles (parameter sets). As the particles move around (i.e., as the algorithm explores the parameter space), an objective function is evaluated at each particle location. Particles communicate with one another to determine which has the lowest objective function value. The objective function for each parameter set was used to identify optimal parameter values. Specifically, we used PSO to minimize the weighted sum of squared residuals (WSSR): 
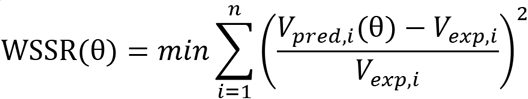
where V*_exp,i_* is the *i*th experimental measurement, V*pred,i* is the *i*th predicted value at the corresponding time point, and *n* is the total number of experimental data points. The minimization is subject to θ, the set of upper and lower bounds on each of the free parameters. The bounds were set to be one order of magnitude above and below the baseline parameter values, which were taken from literature and listed in Tables S3-S4.

The model was fitted against three datasets, specifically: 1) normalized pERK induced by FGF concentrations varying from 0.16 ng/ml to 500 ng/ml, where pERK level was normalized by the maximum pERK stimulated by FGF across all six concentrations (0.16, 0.8, 4, 20, 100, and 500 ng/ml) in two hours, experiments conducted using the H1703 cell line [21]; 2) normalized pVEGFR2 (pR2) stimulated by 5 ng/ml VEGF, pR2 was normalized by the maximum pR2 induced by 5 ng/ml VEGF, experiments conducted using HUVECs [23]; 3) normalized pERK induced by 50 ng/ml VEGF, where pERK was normalized by the maximum pERK induced by 50 ng/ml VEGF, experiments conducted using HUVECs [22]. We note that that the pERK and pR2 in the model simulation include all free and bound forms of pERK and ppERK, and all free and bound forms of pR2 except the degraded pR2, respectively.

After model training, we validated the fitted model with three datasets not used in the fitting. First, we simulated the effects following the addition of heparin. For this case, we added 500 µg/ml heparin, which competes with HSGAG and binds to FGF. There are an additional 26 reactions, 25 species, and 3 parameters for heparin perturbation in the model. Without any fitting, parameters are all taken from Kanodia *et al*. Details are provided in Tables S5-S6. We simulated the pERK dose response with or without heparin to compare with the experiments described by Kanodia [21]. Having a difference in pERK with and without additional heparin that is greater than zero indicates that the presence of heparin enhances FGF-induced ERK phosphorylation. Second, we predicted the phosphorylated pERK following stimulation with 10 ng/ml FGF, mimicking measurements obtained from BAECs [27]. Third, we predicted the VEGF-induced pR2 response upon stimulation with 80 ng/ml VEGF, simulating experiments conducted using HUVECs [28]. For all three datasets, we simulated the experimental conditions without any additional model fitting and compared to the experimental measurements. A total of 16 parameter sets with the smallest errors were taken to be the “best” sets based on the model fitting and validation (Figure S1 and Table S1) and were used for model simulations.

*VEGFR2 density*. To study the impact of VEGFR2 expression on VEGF-induced angiogenesis, we varied VEGFR2 density within ten-fold of the baseline value (1,000 molecules/cell) and predicted the level and dynamics of ERK phosphorylation.

*VEGFR2 trafficking parameters*. To investigate the effects of VEGFR2 trafficking in VEGF-induced ERK phosphorylation, we decreased the trafficking parameters (internalization, recycling, and degradation rates) values for VEGFR2. We changed the parameters individually or together to be the same level as FGFR trafficking parameters and predicted the VEGF-induced pERK response.

### ERK phosphorylation response

We investigated the ERK phosphorylation response by the stimulation of FGF or VEGF alone, compared to their combination. In this study, we mainly focus on two aspects of pERK dynamics: magnitude of the response and timescale of signaling.

### Magnitude of ERK phosphorylation response

a. *Maximum pERK*. We calculate the maximum ERK phosphorylation level induced by the stimulation of FGF, VEGF, or their combination.
b. *Ratio, R*. To compare the combination effects with FGF and VEGF individual effects, we introduce the ratio below: 
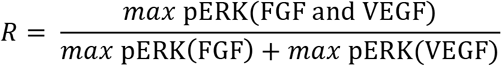
When *R* is greater than one, it indicates that the combination effect in inducing maximal pERK is greater than the summation of individual effects; when *R* is equal to one, it implies that the combination effect is additive; when *R* is less to one, it suggests an antagonistic effect between FGF and VEGF.

*Timescale of the signaling response*. We use two parameters to characterize the timescale of ERK activation: the time to reach the maximum pERK (*T1*) and the time duration that pERK level remains greater than half of its maximal value (*T2*). T1 indicates how quickly ERK is phosphorylated: the smaller T1 is, the faster ERK becomes phosphorylated. T2 indicates how long ERK remains in a phosphorylated state: the larger T2 is, the more sustained the pERK response.

*Reaction rates*. We specify the rates of each reaction based on the law of mass action, where the rate of a chemical reaction is proportional to the amount of each reactant. For example, for the phosphorylation of VEGFR2: 
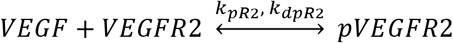

The reaction rate is: 
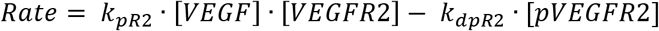
where *k_pR2_* and *k_dpR2_* are rate constants for the forward and reverse reactions, respectively, and [VEGF], [VEGFR2], and [pVEGFR2] are the species’ concentrations.

## Acknowledgements

The authors thank members of the Finley research group for critical comments and suggestions. The authors acknowledge the support of the US National Science Foundation (CAREER Award 1552065). The funders had no role in study design, data collection and analysis, decision to publish, or preparation of the manuscript.

## Additional Files

**Additional File 1 (.xls file):** Supplemental Tables.

**Additional File 2 (.pdf file):** Supplemental Figures.

**Additional File 3 (.docx file):** MATLAB file containing computational model.

## References

1. Loffredo F, Lee RT. Therapeutic vasculogenesis: it takes two. Circ Res. 2008;103(2):128–30.

2. Griffith CK, Miller C, Sainson RC, Calvert JW, Jeon NL, Hughes CC, et al Diffusion limits of an in vitro thick prevascularized tissue. Tissue Eng. 2005;11(1–2):257–66.

3. Lovett M, Lee K, Edwards A, Kaplan DL. Vascularization strategies for tissue engineering. Tissue Eng Part B Rev. 2009;15(3):353–70.

4. Al-Husein B, Abdalla M, Trepte M, Deremer DL, Somanath PR. Antiangiogenic therapy for cancer: an update. Pharmacotherapy. 2012;32(12):1095–111.

5. Keating GM. Bevacizumab: a review of its use in advanced cancer. Drugs. 2014;74(16):1891–925.

6. Vasudev NS, Reynolds AR. Anti-angiogenic therapy for cancer: current progress, unresolved questions and future directions. Angiogenesis. 2014;17(3):471–94.

7. Simons M, Annex BH, Laham RJ, Kleiman N, Henry T, Dauerman H, et al Pharmacological treatment of coronary artery disease with recombinant fibroblast growth factor-2: double-blind, randomized, controlled clinical trial. Circulation. 2002;105(7):788–93.

8. Henry TD, Annex BH, McKendall GR, Azrin MA, Lopez JJ, Giordano FJ, et al The VIVA trial: Vascular endothelial growth factor in Ischemia for Vascular Angiogenesis. Circulation. 2003;107(10):1359–65.

9. Sasich LD, Sukkari SR. The US FDAs withdrawal of the breast cancer indication for Avastin (bevacizumab). Saudi Pharm J. 2012;20(4):381–5.

10. Korc M, Friesel RE. The role of fibroblast growth factors in tumor growth. Curr Cancer Drug Targets. 2009;9(5):639–51.

11. Tan WH, Popel AS, Mac Gabhann F. Computational model of VEGFR2 pathway to ERK activation and modulation through receptor trafficking. Cell Signal. 2013;25(12):2496–510.

12. Asahara T, Bauters C, Zheng LP, Takeshita S, Bunting S, Ferrara N, et al Synergistic effect of vascular endothelial growth factor and basic fibroblast growth factor on angiogenesis in vivo. Circulation. 1995;92(9 Suppl):II365–71.

13. Pepper MS, Ferrara N, Orci L, Montesano R. Potent synergism between vascular endothelial growth factor and basic fibroblast growth factor in the induction of angiogenesis in vitro. Biochem Biophys Res Commun. 1992;189(2):824–31.

14. Mac Gabhann F, Popel AS. Targeting neuropilin-1 to inhibit VEGF signaling in cancer: Comparison of therapeutic approaches. PLoS Comput Biol. 2006;2(12):e180.

15. Stefanini MO, Wu FT, Mac Gabhann F, Popel AS. Increase of plasma VEGF after intravenous administration of bevacizumab is predicted by a pharmacokinetic model. Cancer Res. 2010;70(23):9886–94.

16. Filion RJ, Popel AS. Intracoronary administration of FGF-2: a computational model of myocardial deposition and retention. Am J Physiol Heart Circ Physiol. 2005;288(1):H263–79.

17. Wu Q, Finley SD. Predictive model identifies strategies to enhance TSP1-mediated apoptosis signaling. Cell Commun Signal. 2017;15(1):53.

18. Zheng XM, Koh GY, Jackson T. A Continuous Model of Angiogenesis: Initiation, Extension, and Maturation of New Blood Vessels Modulated by Vascular Endothelial Growth Factor, Angiopoietins, Platelet-Derived Growth Factor-B, and Pericytes. Discrete Cont Dyn-B. 2013;18(4):1109–54.

19. Chambard JC, Lefloch R, Pouyssegur J, Lenormand P. ERK implication in cell cycle regulation. Biochim Biophys Acta. 2007;1773(8):1299–310.

20. Lassoued W, Murphy D, Tsai J, Oueslati R, Thurston G, Lee WM. Effect of VEGF and VEGF Trap on vascular endothelial cell signaling in tumors. Cancer Biol Ther. 2010;10(12):1326–33.

21. Kanodia J, Chai D, Vollmer J, Kim J, Raue A, Finn G, et al Deciphering the mechanism behind Fibroblast Growth Factor (FGF) induced biphasic signal-response profiles. Cell Commun Signal. 2014;12:34.

22. Bruns AF, Herbert SP, Odell AF, Jopling HM, Hooper NM, Zachary IC, et al Ligand-stimulated VEGFR2 signaling is regulated by co-ordinated trafficking and proteolysis. Traffic. 2010;11(1):161–74.

23. Lamalice L, Houle F, Huot J. Phosphorylation of Tyr1214 within VEGFR-2 triggers the recruitment of Nck and activation of Fyn leading to SAPK2/p38 activation and endothelial cell migration in response to VEGF. J Biol Chem. 2006;281(45):34009–20.

24. Filion RJ, Popel AS. A reaction-diffusion model of basic fibroblast growth factor interactions with cell surface receptors. Ann Biomed Eng. 2004;32(5):645–63.

25. Dupree MA, Pollack SR, Levine EM, Laurencin CT. Fibroblast growth factor 2 induced proliferation in osteoblasts and bone marrow stromal cells: a whole cell model. Biophys J. 2006;91(8):3097–112.

26. Zhao B, Zhang C, Forsten-Williams K, Zhang J, Fannon M. Endothelial cell capture of heparin-binding growth factors under flow. PLoS Comput Biol. 2010;6(10):e1000971.

27. Rikitake Y, Kawashima S, Yamashita T, Ueyama T, Ishido S, Hotta H, et al Lysophosphatidylcholine inhibits endothelial cell migration and proliferation via inhibition of the extracellular signal-regulated kinase pathway. Arterioscler Thromb Vasc Biol. 2000;20(4):1006–12.

28. Chabot C, Spring K, Gratton JP, Elchebly M, Royal I. New role for the protein tyrosine phosphatase DEP-1 in Akt activation and endothelial cell survival. Mol Cell Biol. 2009;29(1):241–53.

29. Imoukhuede PI, Popel AS. Quantitative fluorescent profiling of VEGFRs reveals tumor cell and endothelial cell heterogeneity in breast cancer xenografts. Cancer Med. 2014;3(2):225–44.

30. Zheng J, Wen Y, Song Y, Wang K, Chen DB, Magness RR. Activation of multiple signaling pathways is critical for fibroblast growth factor 2-and vascular endothelial growth factor-stimulated ovine fetoplacental endothelial cell proliferation. Biol Reprod. 2008;78(1):143–50.

31. Zheng J, Bird IM, Melsaether AN, Magness RR. Activation of the mitogen-activated protein kinase cascade is necessary but not sufficient for basic fibroblast growth factor-and epidermal growth factor-stimulated expression of endothelial nitric oxide synthase in ovine fetoplacental artery endothelial cells. Endocrinology. 1999;140(3):1399–407.

32. Bai Y, Leng Y, Yin G, Pu X, Huang Z, Liao X, et al Effects of combinations of BMP-2 with FGF-2 and/or VEGF on HUVECs angiogenesis in vitro and CAM angiogenesis in vivo. Cell Tissue Res. 2014;356(1):109–21.

33. Olsson AK, Dimberg A, Kreuger J, Claesson-Welsh L. VEGF receptor signalling – in control of vascular function. Nat Rev Mol Cell Biol. 2006;7(5):359–71.

34. Gourlaouen M, Welti JC, Vasudev NS, Reynolds AR. Essential role for endocytosis in the growth factor-stimulated activation of ERK1/2 in endothelial cells. J Biol Chem. 2013;288(11):7467–80.

35. Padera R, Venkataraman G, Berry D, Godavarti R, Sasisekharan R. FGF-2/fibroblast growth factor receptor/heparin-like glycosaminoglycan interactions: a compensation model for FGF-2 signaling. FASEB J. 1999;13(13):1677–87.

36. Tong S, Yuan F. Dose response of angiogenesis to basic fibroblast growth factor in rat corneal pocket assay: II. Numerical simulations. Microvasc Res. 2008;75(1):16–24.

37. Hendrata M, Sudiono J. A Computational Model for Investigating Tumor Apoptosis Induced by Mesenchymal Stem Cell-Derived Secretome. Comput Math Methods Med. 2016;2016:4910603.

38. Kano MR, Morishita Y, Iwata C, Iwasaka S, Watabe T, Ouchi Y, et al VEGF-A and FGF-2 synergistically promote neoangiogenesis through enhancement of endogenous PDGF-BPDGFRbeta signaling. J Cell Sci. 2005;118(Pt 16):3759–68.

39. Murakami M, Nguyen LT, Hatanaka K, Schachterle W, Chen PY, Zhuang ZW, et al FGF-dependent regulation of VEGF receptor 2 expression in mice. J Clin Invest. 2011;121(7):2668–78.

40. Lampugnani MG, Orsenigo F, Gagliani MC, Tacchetti C, Dejana E. Vascular endothelial cadherin controls VEGFR-2 internalization and signaling from intracellular compartments. J Cell Biol. 2006;174(4):593–604.

41. Sperinde GV, Nugent MA. Mechanisms of fibroblast growth factor 2 intracellular processing: a kinetic analysis of the role of heparan sulfate proteoglycans. Biochemistry. 2000;39(13):3788–96.

